# An innovative approach for determining composite wheat quality index to identify quality enriched genotypes - insights and implications

**DOI:** 10.1101/2021.05.15.444291

**Authors:** D Mohan, R Sendhil, Om Prakash Gupta, Vanita Pandey, K Gopalareddy, Gyanendra Pratap Singh

## Abstract

Ranking test entries or test sites on a quality basis is very difficult in wheat as value addition is perceived by several grain properties and end-products. Here, a novel approach has been developed and tested by deriving wheat quality index based on principal component analysis of 13 physico-chemical grain parameters and 3 end products of 45 wheat varieties. Depending upon the observed index range (0.15 to 0.71), the cultivars were assorted into 3 distinct classes *i*.*e*. elite, moderate and poor. The top group ascertained high quality standards of grain suited for bread and *chapati* whereas bottom group assured better cookies quality. This technique was also tested to differentiate quality enriched test sites within a zone or demarcate the most suited production environments to harness good quality wheat. The index will have an implication on farmers (premium price for varietal segregation), industry (product specific quality cultivars), and consumers (superior quality products).

## 1. Introduction

Wheat (*Triticum* spp.) alone constitutes nearly one-third of the world’s total cereal consumption (FAO, 2003) and is a staple food in many countries including India. Although this staple food crop is mostly consumed as unleavened flat bread (*chapati*), 15% of the harvested produce goes to the baking industry of the country for different bakery products including bread and cookies. Value addition properties of wheat therefore, are vital not only for domestic consumption but for the baking industry as well. Wheat quality has many shades and presently, it is described through various parameters. Wheat varieties can easily be distinguished for a single quality parameter. Wheat quality is important for different stakeholders in the wheat value chain i.e. farmers (bold and plump grain), millers (test weight and flour yield), food processors (processing quality) and consumers (end-use and nutritional quality) (Guzman et al., 2019). There are number of component quality parameters including grain appearance, test weight, grain protein, grain hardness, sedimentation, gluten content, gluten index, iron, zinc, phenol score, flour extraction were utilized either individually or in a combination to categorize the wheat varieties suitable for specific end-products like bread, biscuit, and *chapatti* score. Based on these one or two component traits, wheat varieties have been classified in to different product specific genotypes in different countries and HS490 for better biscuit quality is one such example in India.

The relationship of GlutoPeak indices with various conventional quality parameters including grain hardness, sedimentation value, farinograph, alveograph were studied to ascertain the utility of GlutoPeak test to predict the wheat flour baking properties (Gucbilmez et al., 2019). SDS sedimentation test could be utilized to predict the baking quality and gluten strength and also as a rapid test in wheat breeding programs (Carter et al., 1999). Soft endosperm genes in wheat are responsible for better biscuit-making ability through associated traits like low alveograph stability, strength, P/L ratio, and protein content, and high alveograph extensibility, biscuit diameter (Labuschagne et al., 1997). In a similar study to unravel the effect of soft endosperm on biscuit quality was attempted by Ma et.al. (2018) and revealed that the soft wheat varieties with low protein contents (7.9-9.7%), low sedimentation volume (20.0-32.0 ml), and low damaged starch contents (1.9-3.4%) are desirable traits for good biscuit making quality. A study of different physico-chemical parameters including grain appearance score, hardness, test weight, thousand kernel weight, protein, gluten content and index, sedimentation value, phenol test, carotenoids, diastatic activity to ascertain their role in *chapati* making quality by Kumar et al., (2018) revealed a clear role of grain hardness and diastase activity, conversely phenol score may not serve as a suitable indicator of chapatti quality. Bonafede et. al. (2015) studied wheat NILs to understand the effect of *Glu-3* and *Gli-1* loci on bread making quality through different allelic combinations of *Glu3/Gli-1*.

Genetic resource good for respective quality trait had been distinguished from the commercial wheat varieties in India too (Mohan et al.,, 2013), but identification based on multiple quality traits is rather complex and has not been attempted yet. It always desired that the varieties picked for a particular end-product should also possess a combination of other desirable attributes as well like grain appearance, nutritional values, processing quality etc. A variety with high baking potential and good a combination of other desired grain quality attributes is certainly better than the one which has good end-product quality but lacks in some important grain properties. There are also incidents when a genotype possesses a good combination of the desired grain quality features but the quality of the end-products is not up to the mark. At a time when the relevance of good quality wheat is picking up across the globe, a uniform system is utmost important to differentiate the high-rank wheat cultivars with several good quality features. Although it sounds astonishing to find a genotype which has all desired value addition properties but the varietal distinction is necessary especially when a big bunch of cultivars is under cultivation. This study is an attempt to address this snag by converting the multivariate wheat quality evaluation into a single window system named as ‘Wheat Quality Index’. The data generated by the All India Coordinated Research Project on Wheat and Barley (AICRPW&B) on grain quality attributes of popular irrigated bread wheat varieties have been used to test this unique statistical approach which can be useful to rank value addition properties of wheat germplasm.

## 2. Materials and methods

### 2.1 Study materials and value addition properties

Popular wheat varieties used as a check in timely-sown (TS) and late-sown (LS) yield evaluation trial series conducted by AICRPW&B in five mega zones of the country *i.e*. Northern Hills Zone (NHZ), North Western Plains Zone (NWPZ), North Eastern Plains Zone (NEPZ), Central Zone (CZ) and Peninsular Zone (PZ); were selected for this investigation. The study material involved four years of performance (period: 2014-17) of 45 high yielding irrigated wheat varieties at 3-5 test sites. Grain samples received from each test site were analyzed at the headquarter of ICAR-IIWBR located at Karnal, India as per the international standards (AACC, 2000). Data recorded on 13 grain quality parameters includes grain appearance score, test weight, grain hardness index, sedimentation value, grain protein content at 14% grain moisture, grain load on protein content *i.e*. protein yield per hectare, wet gluten content, gluten index, *Glu 1* score, iron and zinc contents and extraction rate. Besides, phenol test score data was also included as polyphenol oxidase (PPO) activity is known to have strong negative impact on *chapati* quality. Data about end-products quality included *chapati* quality score, bread loaf volume, bread quality score and biscuit spread factor.

### 2.2 Statistical analysis

Four year’s mean value of each variety for each parameter were computed and principal component analysis (PCA) was used to derive the composite quality index based upon a whole array of grain quality attributes. Since the wheat quality indicator variables have different units of measurement (see Table S1, column 1), they were normalized using the following formula (Eq. 1 and 2) to make them scale-free for comparison as outlined by Mahida and Sendhil (2017). Equation (1) shows the normalization formula used for indicator variables having a positive functional relationship with the wheat quality as observed in bread and *chapati*.

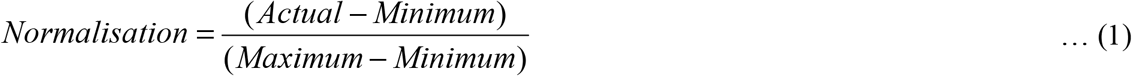

In the case of variables that have a negative association with end-products like biscuit spread factor, the following equation (2) was used.

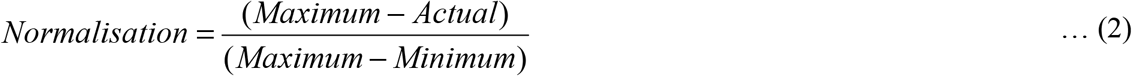

Post normalization, weights have to be assigned to the selected wheat quality indicator variables so that aggregation can be done to derive the composite WQI. We adopted the PCA approach for the calculation of weights due to its merit over other available techniques as well as based on Kaiser criterion (Kaiser, 1960) that selects the principal components having more than ‘one’ Eigen value to capture the maximum variation in the data matrix. We used the functional framework as indicated in Equation (3).

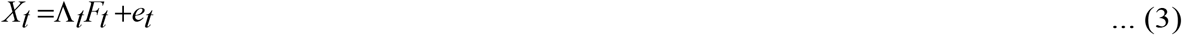

where, X_t_ is the N-dimensional vector of variables affecting the wheat quality, Λ*t* is rx1 common factor, *F*_*t*_ is the factor loading, and *e*_*t*_ is the associated idiosyncratic error-term of order Nx1. Weights for each wheat quality variable were calculated from the PCA results as indicated in Equation (4)

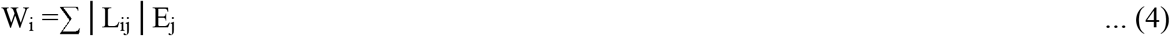

where, W_i_ is the weight of the i^th^ indicator variable, E_j_ is the Eigen value of the j^th^ factor, and L_ij_ is the loading value of the i^th^ variable on j^th^ factor. After deriving the weights for all the variables, the composite WQI for each genotype has been calculated using Equation (5)

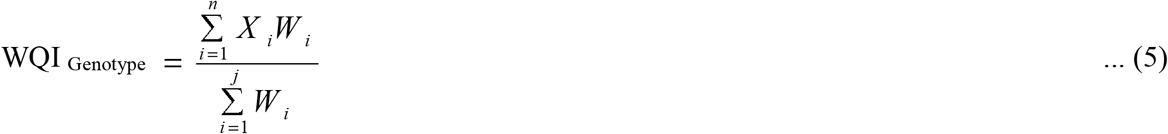

where X_i_ is the normalized value of i^th^ wheat quality indicator variable and W_i_ is the weight of the i^th^ variable. This technique had been earlier used by Mamrutha et al., (2020) to develop stress screening index for prioritizing hotspot locations for wheat under Indian environments and Rana et al., (2015) to derive salt tolerance index. Based upon WQI, the genotypes were classified into three categories *i.e*. elite, moderate and poor by using the following formula.

- Elite = WQI> (Mean + 0.5 Standard deviation)
- Moderate = (Mean – 0.5 Standard deviation) <WQI< (Mean + 0.5 Standard deviation)
- Poor = WQI< (Mean – 0.5 Standard deviation)

The quality index was derived for bread and *chapati* separately and in combinations as well. Barring phenol test score, the preference to wheat grain quality was positive in all such computations. Trait specificity was just the opposite, while deriving the quality index for the biscuit. Apart from the aforementioned analytical tools and techniques, Pearson’s correlation coefficient was calculated to study the inter-trait relationship and Student’s “t-test” was applied to differentiate grain quality characteristics of two groups. Coefficient of variation (CV) was also calculated to gauge diversity in the study material.

## 3. Results

### 3.1 Diversity in wheat quality parameters

The study material which was a bunch of high yielding popular wheat varieties of diverse Indian production environments, expressed diversity in several grain quality attributes (Table S1). Since the principal component analysis is largely based upon the variation level and inter-trait relationship; extent of variation was examined in the study material. Diversity level was very high in phenol test score, gluten index, sedimentation value and protein yield (CV: above 15 %); moderate in biscuit spread factor, grain hardness index, wet gluten content, and Glu 1 score (CV: 10 to 15%); low in bread quality score, grain appearance score, iron, zinc and protein contents (CV: 5 to 10%); and very low in bread loaf volume, flour extraction rate, chapatti quality score and test weight (CV: below 5%).

Physical grain quality was good in the majority of the varieties as grain appearance score was ≥ 6.0 in 30 genotypes. Test weight below 78.0 kg/hl was also noticeable only in 9 genotypes. Protein concentration was also satisfactory in the study material as only 10 genotypes expressed protein content below 11.0% whereas protein yield was below 500 kg/ha only in 16 varieties. Since Indian wheats generally have hard grain texture, soft grain texture was witnessed only in one wheat variety cultivated in the hills *i.e*. HS 490. Gluten strength was quite good in the study material as 18 varieties registered sedimentation value ≥ 50ml and just 5 were below 40ml. Overall flour recovery was also satisfactory as extraction rate was ≥ 70% in 19 varieties and below 69% only in 7 genotypes. Reaction to phenol was quite diverse but score below 5.0 was witnessed only in 11 test entries. Gluten content in the selected gene pool was also satisfactory as only 7 genotypes had wet gluten content below 25%. Varieties with a wet gluten content of the range 32-36% were also cited in the study material. Quality of the gluten was also quite befitting as gluten index was below 50% only in 6 genotypes. *Glu 1* score was perfect 10 in 16 genotypes and only 4 genotypes had *Glu 1* score below 8. *Chapati* quality was good in the majority of the material as a score below 7.00 was noticed only in 2 cultivars. Traditionally, bread quality in the commercial Indian wheat cultivars is not rated very high. In this investigation also, only one-third material expressed bread loaf volume ≥ 575cc but the bread quality score over 7.5 was observed only in 4 test entries. Obviously, biscuit quality was not good in this set of material for want of grain softness. Biscuit spread factor ≥ 10.0 was witnessed in the lone soft grain variety *i.e*. HS 490 but biscuit spread factor above 8.5 could be witnessed in two hard grain varieties of NHZ.

### 3.2 Germplasm assortment for the end-products

#### 3.2.1 Quality index for bread and chapati

This holistic approach widened the difference between 45 test entries as the quality index varied from 0.197 to 0.717 in *chapati* and 0.165 to 0.724 in bread. In the elite groups of bread and *chapati*, 15 varieties occupied each cluster out of which 13 were superior for both bread as well as *chapati* (Table 1). Since the majority of the varieties were common in the two clusters, another attempt was made to derive quality index which involved *chapati* as well bread quality parameters and it was named as the wheat quality index (WQI). Range of WQI was almost similar (0.154 to 0.711) to quality index of bread or *chapati*. The elite group formed on this basis also contained 15 genotypes and this cluster involved all 15 varieties of the elite bread group and 13 entries of the elite *chapati* group. Bread and *chapati* quality in the high ranking genotypes was quite good as cluster mean was 573cc for bread loaf volume and 7.77 for *chapati* quality score.

**Table 1:** Elite varieties for the quality index of bread and *chapati*

| Variety | Production environment | Bread | *Chapati* | Bread cum *chapati* |
| --- | --- | --- | --- | --- |
| RAJ 4083 | PZ-LS | 0.724 | 0.717 | 0.711 |
| HD 2932 | PZ-LS | 0.696 | 0.701 | 0.706 |
| UAS 304 | PZ-TS | 0.676 | 0.637 | 0.675 |
| HD 3226 | NWPZ-TS | 0.661 | 0.644 | 0.648 |
| PBW 752 | NWPZ-LS | 0.651 | 0.658 | 0.632 |
| NIAW 917 | PZ-TS | 0.650 | - | 0.632 |
| DBW 90 | NWPZ-LS | 0.627 | 0.654 | 0.622 |
| HD 3059 | NWPZ-LS | 0.626 | 0.651 | 0.623 |
| HD 2864 | CZ-LS | 0.618 | 0.647 | 0.634 |
| MP 4010 | CZ-LS | 0.615 | 0.632 | 0.610 |
| MP 3336 | CZ-LS | 0.615 | 0.657 | 0.629 |
| HD 3090 | PZ-LS | 0.612 | - | 0.599 |
| MACS 6478 | PZ-TS | 0.612 | 0.633 | 0.636 |
| WH 1124 | NWPZ-LS | 0.601 | 0.643 | 0.604 |
| HD 2932 | CZ-LS | 0.606 | 0.630 | 0.622 |
| DBW 173 | NWPZ-LS | - | 0.605 | - |
| HD 3249 | NEPZ-TS | - | 0.602 | - |

#### 3.2.2 Quality index for cookies

In this approach, bread and *chapati* parameters were excluded and the quality index was derived with biscuit spread factor. Variation was quite large 0.196 to 0.702 for biscuit quality index also. Again, 15 varieties occupied the top group in biscuit quality index and the average biscuit spread factor was 8.0 (Table S2). There is only one soft wheat variety in the country which has biscuit spread factor more than 10 *i.e*. HS 490 and this genotype occupied the 1^st^ ranking in biscuit quality index (Anonymous, 2016). Although the varieties under study lacked grain softness, all genotypes with biscuit spread factor over 8.0 were noticeable in this elite group. Six varieties in this group had biscuit spread factor below 7.5 but they could find a place in the top group because of some grain properties suited for cookies. The genotype (HS490) ranked 1^st^ in this analysis (HS 490) was the poorest in bread and *chapati* quality indices. In fact, a majority of the entries clustered in the poor group, were the ones which were placed in the elite group formed based on bread or *chapati* alone.

Three varieties each of NWPZ (HD 3086, WH 1105 and PBW 550) and NEPZ (DBW 39, NW 2036 and HI 1563) registered moderate quality standards in all the products. There was no wheat variety which could be rated inferior for every product. There were certain exceptional cases where the product quality was good but the quality index was low. It happened in the case of HD 2733 for bread (loaf volume: 585cc) and DBW 168 for *chapati* (score: 8.15). HD 2733 suffered because of grain protein content and gluten properties whereas DBW 168 lacked in grain hardness and extraction rate.

#### 3.2.3 Screening advantage with wheat quality index

Since a majority of the varieties placed in the bread and *chapati* indexes featured in the combined analysis, WQI was further examined for usefulness in germplasm assortment. This index alone can also be applied to screen the biscuit quality enriched germplasm as well. To substantiate it further, the elite and poor groups formed on the basis WQI were compared for the end-products quality (Table 2). Bread or *chapati* quality in the elite group was significantly better than the poor group. On the contrary, biscuit spread factor in the poor group was significantly higher than the elite group. It reconfirmed that WQI alone can be quite useful for varietal distinction in different end-products.

**Table 2:**
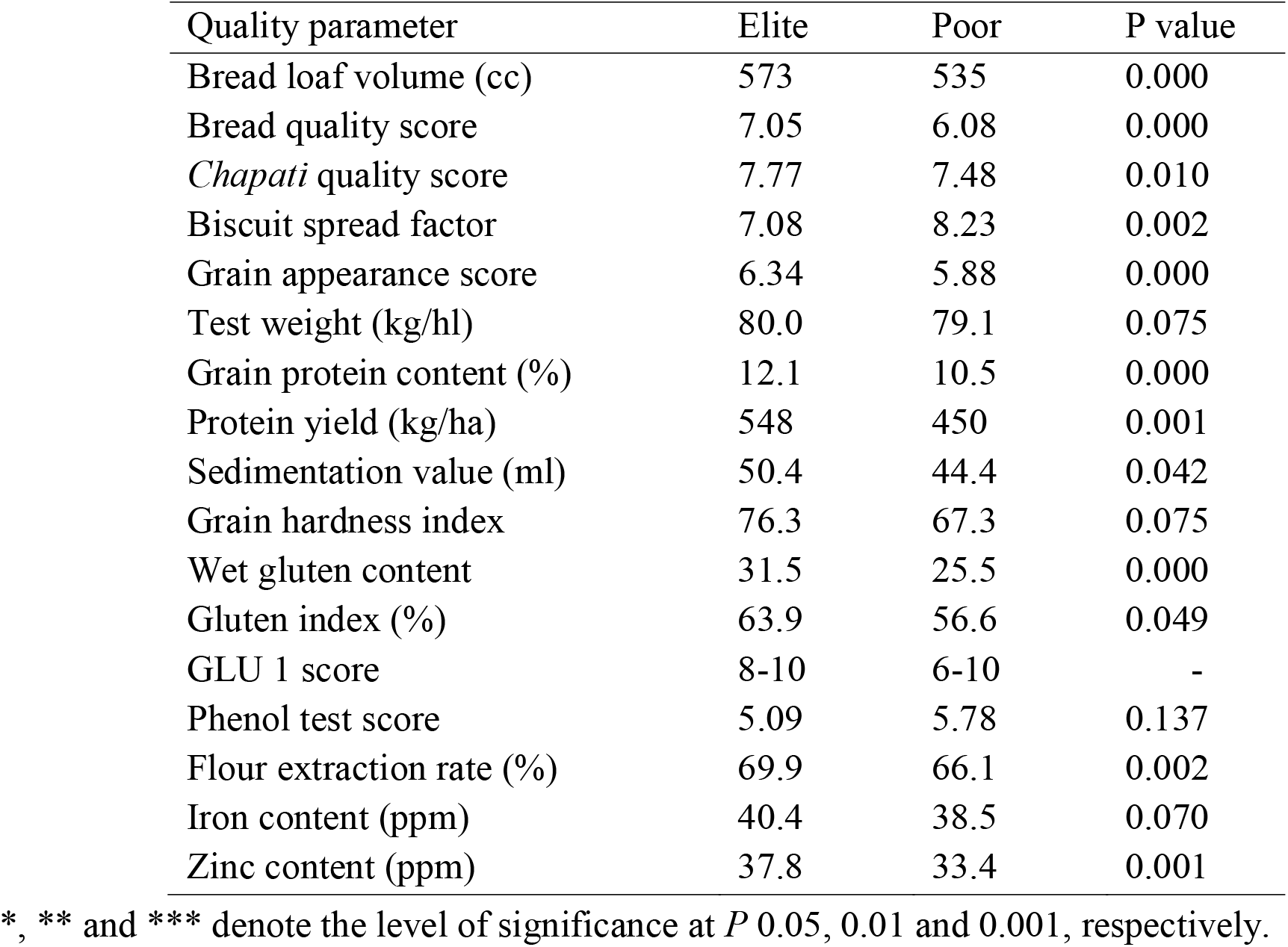
Mean values of grain quality parameter in diverse quality groups

Besides bread and *chapati* quality, the elite material was superior in grain appearance, protein content and protein yield, gluten properties like gluten content, gluten strength (sedimentation value) and gluten quality (gluten index), flour recovery and grain zinc density. The difference in grain hardness could not be a witness and it was not expected also as the study material was largely devoid of soft grains. A significant difference in the two contrasting group could not be established in test weight, iron content and phenol reaction. It means that test weight, grain hardness, iron and phenol reaction did not play any decisive role in defining WQI of the Indian varieties involved in this study.

#### 3.2.4 Quality features in the screened elite material

The quality index in the elite material screened based on WQI ranged from 0.60 to 0.71. In *chapati* quality, genotypes with score 7.5 to 8.0 are rated very high and every entry of this group had *chapati* score within this range (Table 3). Regression coefficient derived by computing 13 grain quality components with the end-products was found highly significant (*P* 0.001) in *chapati* (R^2^: 0.88). Therefore, prospects of tracing varieties good for *chapati* making were quite high by this technique. Some varieties with *chapati* score around 8.0 or above did occur in the moderate and poor groups also like HI 1563 and DBW 168 but they missed few vital grain quality attributes. These two varieties were just moderate in grain appearance score, test weight and wet gluten content. Protein yield was very low in HI 1563 whereas DBW 168 lacked in grain hardness and gluten index.

**Table 3:** Value addition properties of elite varieties for good *chapati* and bread quality

| Variety | Environment | GAS | TW | GPC | Fe | Zn | SV | Glu | GHI | FER | WGC | GI | CQS | BLV | BQS | BSF | Grade |
| --- | --- | --- | --- | --- | --- | --- | --- | --- | --- | --- | --- | --- | --- | --- | --- | --- | --- |
| **I. Good quality genotypes** |  |  |  |  |  |  |  |  |  |  |  |  |  |  |  |  |  |
| HD 2932 | PZ-LS | 6.3 | 80 | 12.7 | 40 | 38 | 47 | 8 | 72 | 70.2 | 36 | 62 | 8.0 | 585 | 7.5 | 7.4 | A<sup>++</sup> |
| UAS 304 | PZ-TS | 6.3 | 81 | 11.7 | 47 | 43 | 44 | 8 | 75 | 69.9 | 32 | 49 | 7.8 | 595 | 7.9 | 6.7 | A<sup>+</sup> |
| NIAW 917 | PZ-TS | 6.1 | 82 | 11.8 | 46 | 36 | 41 | 8 | 84 | 70.0 | 31 | 61 | 7.5 | 590 | 7.9 | 6.7 | A<sup>+</sup> |
| RAJ 4083 | PZ-LS | 6.6 | 80 | 12.1 | 43 | 40 | 51 | 10 | 80 | 71.7 | 33 | 68 | 7.7 | 583 | 7.4 | 6.7 | A |
| HD 3090 | PZ-LS | 6.4 | 80 | 12.3 | 42 | 41 | 43 | 8 | 77 | 69.7 | 30 | 60 | 7.6 | 577 | 7.2 | 7.3 | A |
| MACS 6478 | PZ-TS | 6.2 | 79 | 12.0 | 36 | 36 | 50 | 8 | 74 | 70.2 | 33 | 53 | 8.1 | 578 | 7.1 | 6.4 | A<sup>+</sup> |
| HD 2864 | CZ-LS | 6.4 | 82 | 12.1 | 39 | 35 | 45 | 8 | 71 | 69.4 | 31 | 62 | 8.0 | 557 | 6.8 | 7.1 | A<sup>+</sup> |
| MP 3336 | CZ-LS | 6.7 | 81 | 12.6 | 40 | 38 | 39 | 8 | 71 | 69.6 | 34 | 48 | 8.0 | 549 | 6.4 | 7.0 | A<sup>+</sup> |
| HD 2932 | CZ-LS | 6.4 | 81 | 12.6 | 37 | 34 | 49 | 8 | 68 | 69.0 | 33 | 55 | 8.0 | 563 | 7.0 | 6.9 | A<sup>+</sup> |
| MP 4010 | CZ-LS | 6.8 | 82 | 12.7 | 40 | 38 | 41 | 8 | 70 | 69.2 | 33 | 51 | 7.8 | 556 | 6.5 | 6.8 | A |
| HD 3226 | NWPZ-TS | 6.1 | 78 | 12.0 | 39 | 37 | 64 | 10 | 76 | 69.0 | 301 | 79 | 7.5 | 601 | 7.4 | 7.6 | A |
| PBW 752 | NWPZ-LS | 6.0 | 79 | 11.9 | 40 | 40 | 66 | 10 | 83 | 69.5 | 29 | 83 | 7.5 | 577 | 6.9 | 7.0 | A |
| HD 3059 | NWPZ-LS | 6.3 | 79 | 11.8 | 38 | 37 | 61 | 10 | 82 | 70.2 | 29 | 77 | 7.7 | 569 | 6.7 | 7.6 | A |
| DBW 90 | NWPZ-LS | 6.2 | 78 | 11.5 | 41 | 37 | 60 | 10 | 83 | 70.5 | 29 | 76 | 7.6 | 565 | 6.6 | 7.5 | A |
| WH 1124 | NWPZ-LS | 6.3 | 78 | 11.2 | 40 | 39 | 58 | 10 | 81 | 70.4 | 28 | 77 | 7.7 | 556 | 6.5 | 7.5 | A |
| **II. Poor quality genotypes** |  |  |  |  |  |  |  |  |  |  |  |  |  |  |  |  |  |
| DBW 168 | PZ-TS | 5.9 | 78 | 11.7 | 38 | 34 | 43 | 6 | 36 | 67.0 | 29 | 44 | 8.2 | 555 | 6.7 | 7.9 | C<sup>+</sup> |
| GW 322 | PZ-TS | 6.0 | 79 | 11.0 | 36 | 39 | 40 | 8 | 80 | 70.1 | 30 | 51 | 7.7 | 541 | 6.5 | 7.1 | C |
| K 0307 | NEPZ-TS | 6.1 | 79 | 11.2 | 42 | 36 | 40 | 8 | 79 | 70.7 | 27 | 57 | 7.6 | 536 | 6.0 | 7.8 | C |
| DBW 187 | NEPZ-TS | 5.9 | 78 | 10.6 | 43 | 27 | 65 | 10 | 71 | 69.1 | 26 | 63 | 7.7 | 530 | 5.8 | 8.0 | C |
| K 1006 | NEPZ-TS | 5.9 | 79 | 10.9 | 41 | 36 | 39 | 8 | 78 | 69.9 | 28 | 47 | 7.5 | 557 | 6.4 | 7.9 | C |
| HPW 349 | NHZ-TS | 5.7 | 81 | 9.3 | 35 | 32 | 52 | 10 | 71 | 64.1 | 21 | 80 | 7.4 | 545 | 6.3 | 8.8 | C<sup>+</sup> |
| HS 507 | NHZ-TS | 6.2 | 81 | 9.3 | 39 | 32 | 46 | 10 | 79 | 64.7 | 23 | 61 | 7.4 | 540 | 6.3 | 7.9 | C |
| VL 804 | NHZ-TS | 6.1 | 81 | 10.2 | 33 | 29 | 40 | 6 | 81 | 65.0 | 24 | 62 | 7.7 | 542 | 6.3 | 7.6 | C |
| VL 907 | NHZ-TS | 5.7 | 79 | 9.8 | 36 | 32 | 39 | 8 | 69 | 63.4 | 25 | 52 | 7.3 | 547 | 6.4 | 8.2 | C |
| VL 892 | NHZ-LS | 5.9 | 79 | 11.1 | 41 | 36 | 45 | 8 | 71 | 63.1 | 25 | 52 | 7.0 | 499 | 5.1 | 8.3 | C |
| HS 490 | NHZ-LS | 5.4 | 75 | 10.9 | 39 | 34.9 | 40 | 8 | 26 | 60.6 | 22 | 54 | 6.9 | 497 | 5.0 | 11.1 | C<sup>+</sup> |

**Table 4:**
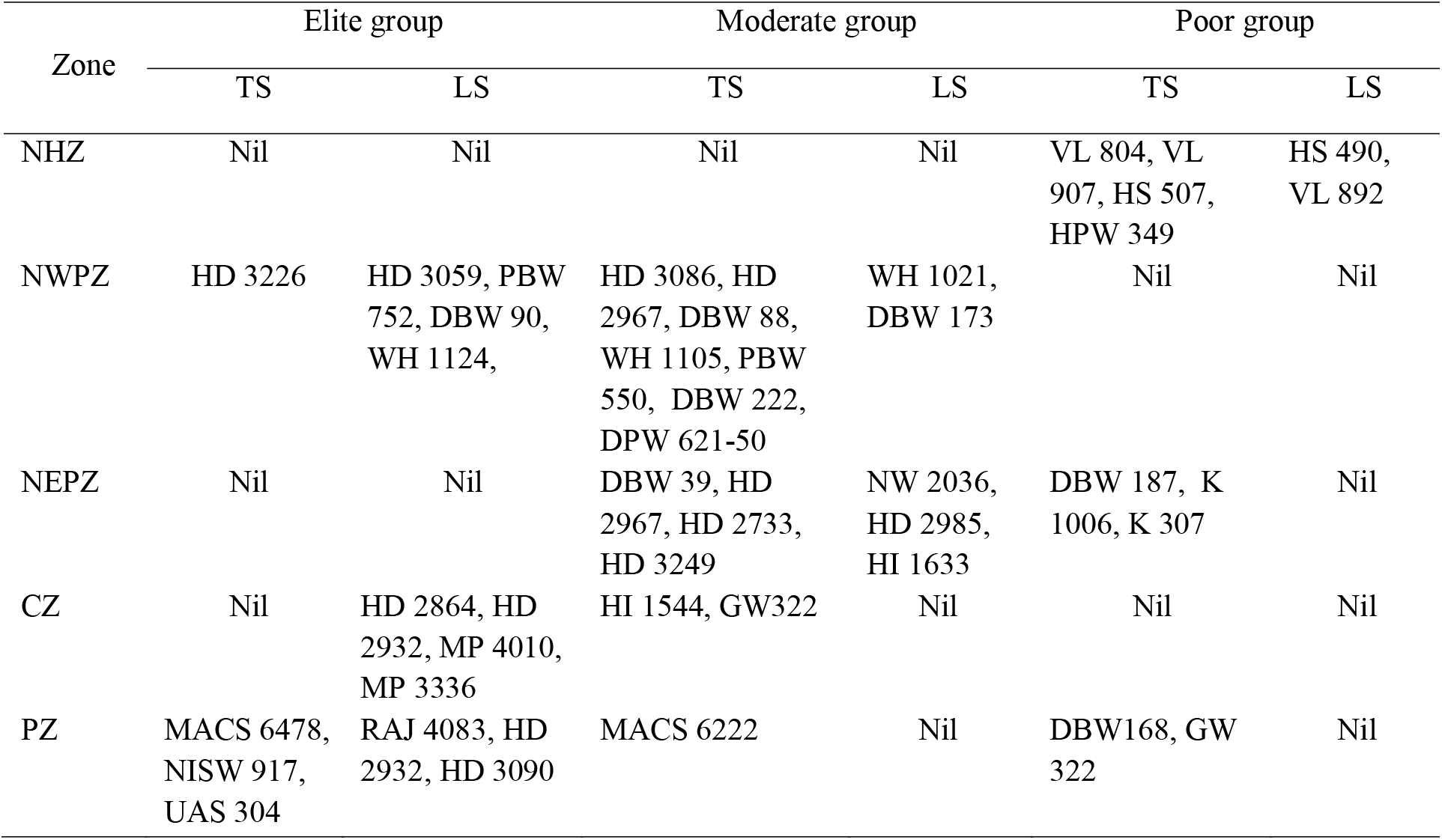
Categorization of Indian wheat varieties in terms of quality and production condition

| Zone | Elite group |  | Moderate group |  | Poor group |  |
| --- | --- | --- | --- | --- | --- | --- |
|  | TS | LS | TS | LS | TS | LS |
| NHZ | Nil | Nil | Nil | Nil | VL 804, VL 907, HS 507, HPW 349 | HS 490, VL 892 |
| NWPZ | HD 3226 | HD 3059, PBW 752, DBW 90, WH 1124, | HD 3086, HD 2967, DBW 88, WH 1105, PBW 550, DBW 222, DPW 621-50 | WH 1021, DBW 173 | Nil | Nil |
| NEPZ | Nil | Nil | DBW 39, HD 2967, HD 2733, HD 3249 | NW 2036, HD 2985, HI 1633 | DBW 187, K 1006, K 307 | Nil |
| CZ | Nil | HD 2864, HD 2932, MP 4010, MP 3336 | HI 1544, GW322 | Nil | Nil | Nil |
| PZ | MACS 6478, NISW 917, UAS 304 | RAJ 4083, HD 2932, HD 3090 | MACS 6222 | Nil | DBW168, GW 322 | Nil |

Quality of the bread is assessed by bread loaf volume and bread quality score. A total of 8 varieties registered bread loaf volume and bread quality score better than mean of the elite cluster. Under Indian condition, wheat varieties with bread loaf volume ≥ 575cc and bread quality score ≥ 7.5 are rated good and varieties like HD 2932, UAS 304 and NIAW 917 qualified this bench mark, too. Regression coefficient achieved by regressing 13-grain quality attributes with this end-product was also highly significant but the magnitude of association was not as high as noticed in the case of *chapati*. In this exercise, R^2^ value was just 0.545 (*P*: 0.008) in case of bread loaf volume and 0.576 (*P*: 0.004) in case of bread quality score. Therefore, the number of high rated bread quality entries was less in the elite group. But when the comparison was made with the poor group; no variety could cross the threshold of 575cc loaf volume and 7.0 bread quality score. Maximum loaf volume and bread quality score achieved in this group were 557cc and 6.71, respectively. It was obvious that good quality bread cannot be made from varieties rated poor by WQI. *Chapati* quality was definitely good in DBW 168 but it could not touch the elite group because of soft grain texture, high bran content and poor gluten quality.

Although the grain quality properties declined in the poor group, chances of locating the good quality cookies were quite high. As done for bread and *chapati*, R^2^ value was calculated for biscuit also by regressing the 13-grain quality parameters against the biscuit spread factor. The regression coefficient was very high in the cookies also (0.83). Therefore, the prospects of discovering cookie material brightened in the cluster where grain quality feature was poor. It underlined that although grain softness is important for biscuit making, it is not absolute and a few other traits also matter in the quality of this product. Chances of locating varieties good for cookies become higher when WQI is applied in the negative direction. All varieties with biscuit spread factor ≥ 8.0 could be seen in the poor group formed based on WQI. At a time when the only soft grain variety placed lowest at WQI ranking *i.e*. HS 490 recorded biscuit spread factor ≥ 10, there was another hard grain variety (hardness index: 71) close to this benchmark *i.e*. HPW 349 with biscuit spread factor 8.82.

### 3.3 Marking regional specificity in value addition

Study material in this analysis belonged to 10 diverse production environments of India. It was evident that material clustered in different groups (Table 1, S2 & 3) did not represent every environment. Grouping made based on WQI indicated that value addition properties in NHZ were poor and the material suited better for cookies. High yielding varieties of NWPZ were either moderate or good in wheat quality. The timely-sown material of this region was generally moderate whereas late-sown varieties excelled in value addition properties. None of the NEPZ varieties could be rated high in wheat quality and they either belonged to moderate or poor category. In this zone, the late-sown wheat was of moderate quality whereas timely-sown wheat could either be moderate or poor. In central India, the late-sown varieties were good in quality whereas timely-sown materials were just moderate. Varieties of the peninsular region were generally good. Couples of varieties like GW 322 and DBW 168 could be rated poor but their *chapati* quality score was quite good.

In India, there are two categories of irrigated wheat *i.e*. timely-sown wheat and late-sown wheat. This technique was applied to know whether these two categories of wheat also express any major difference overall wheat quality. Across the country, 11 varieties figured in the late-sown group whether the number of timely-sown varieties was limited to just 4. In the moderate group however, the number of timely-sown varieties (15) was much higher than the late-sown varieties.

### 3.4 Location specificity within the region

This novel approach based on multivariate grain quality analysis can also be applied to identify the pockets where grain quality can be harnessed better within a given region. Demarcation to verify quality enriched sites also becomes very difficult when the number of variables is high. This multivariate approach was applied to resolve such problems in NWPZ which is not only the most productive wheat land of the country but both production conditions are relevant due to rice-wheat cropping system (Mohan et al.,, 2017). Therefore, it is imperative to decide locations and the appropriate production condition to raise good quality wheat in this high-yield territory. Quality data in this zone was derived from five test sites *i.e*. Ludhiana, Durgapura, Delhi, Hisar and Pantnagar in two production conditions *i.e*. timely-sown and late-sown. Five years mean for the period 2013 to 2017 was computed in each situation and the varieties considered were WH 1105, HD 2967, HD 3059 and DBW 88 for timely-sown wheat and WH 1021, WH 1024, HD 3059 and DBW 90 as late-sown wheat. The quality index, in this case was calculated based on all value addition properties except phenol test score as it did not show any association with the quality index (Table 2). In this exercise, biscuit spread factor was also included and it was the only factor labelled with negative impact in this analysis. Range of WQI for the test centers was also quite large (0.31 to 0.64) therefore it became very easy not only to distinguish good quality sites from the bad ones but also to prioritize the production condition to harness high quality of wheat (Fig 1). Delhi and Durgapur emerged as the pockets best suited to grow high quality wheat in the zone. Results indicated that preference to late-sown wheat at Delhi and timely-sown at Durgapura were the two best options to harness good quality of the harvested produce in NWPZ.

**Fig. 1:**
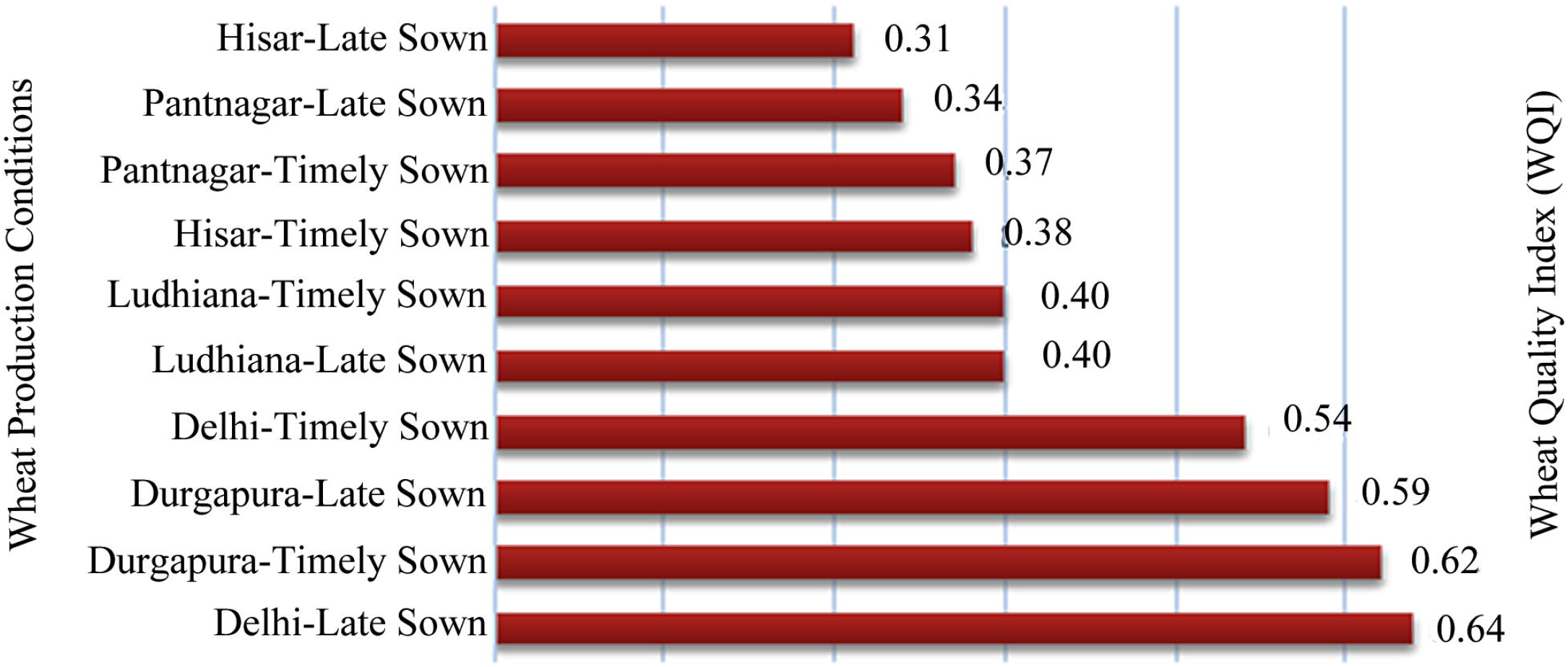
Site selection through the wheat quality index in the north-western plains zone of India. Number indicates wheat quality index for each test center.

## 4. Discussion

Quality of wheat variety cannot be defined by the end-products alone nor wise to look only through the prism of end-products. Wheat quality is multidimensional and cannot be described through a single parameter. But to pick the best material or test site on overall superiority, a single window system is required. This exercise provides a tool where all features of value additions are depicted by a single a factor. And this single factor has been derived by taking into account the overall variation and inter-trait relationship in all wheat quality parameters. This novel and holistic approach is quite sound, flexible and can be applied with any number of variables irrespective of the population size. This study done in the Indian environments involved 17 parameters and the population size was also not big. If the traits of importance are less in some countries or the germplasm under screening is large, even then this technique can be perfected with equal precision. The authors used principal component analysis to put weight to the different quality traits. But this weight can also be assigned depending upon the importance a country or system gives to a particular trait.

### 4.1 The Validity of the wheat quality index

The soundness of this technique was substantiated when value addition properties were examined in the elite and poor groups separated by this method (Table 1). The elite group was significantly better (*P* 0.01) than the poor group in bread and *chapati* quality. It was quite obvious as a strong positive association between *chapati* and bread quality reported earlier by Mohan and Gupta (2008) was reconfirmed in this investigation as well (r: 0.41 ^**^). Contributing traits in bread and *chapati* quality are most common and in contrast to the biscuit (Pena et al.,, 2012; Mohan et al.,, 2013) but there can be genotypes which are good in *chapati* but just moderate in bread making. This technique had also illustrated that genotypes with superior biscuit quality were quite frequent in the cluster where bread or *chapati* quality was generally poor. The genotype identified best through this technique *i.e*. HD 2932 was superior most in bread and *chapati* qualities but poorest in biscuit making. Similarly, the genotypes identified best in biscuit quality index *i.e*. HS 490 had highly inferior quality of bread and *chapati*. It was quite obvious and this well-established fact was again confirmed when the adverse relationship of this product was observed between bread (r: -0.40 ^**^) and *chapati* (r: -0.55 ^**^) in this bunch of high-yield genotypes. In Indian wheat, value addition property of late-sown wheat is rated better than the timely-sown (Mohan et al.,, 2011, 2017) and this technique also supported this trend. All these observations underline that this technique holds true and is quite sound in judging overall the quality of wheat.

### 4.2 Usefulness

This technique discriminates the material where product quality is good but remains unsupported by the grain quality determinants as observed in the case of DBW 168 for *chapati* and HD 2733 for bread loaf volume (Table 3). Despite of good product quality, none of them could occupy place in the elite group. On the contrary, it does elate the materials which have several product supporting grain quality features but the product quality is somehow not so good. In wheat breeding, selection strategy for quality improvement differs product-wise (Mohan et al., 2013) but perfection about the roles that grain quality parameters play in defining the product quality, is yet to be achieved. This novel approach which offers best a genotype can provide in value addition is the best option for each end-product (Fig 2). When applied in the positive direction it supports bread or *chapati* making and in reverse, quality of the cookies is improvised. India has the second-largest biscuit industry in the world which cannot depend only on single soft grain genotype. Definitely, this industry must not have survived or it cannot survive based on one genotype. This analysis supports that when soft wheat is rarely cultivated, there can also be other alternates to support this industry.

**Fig. 2.**
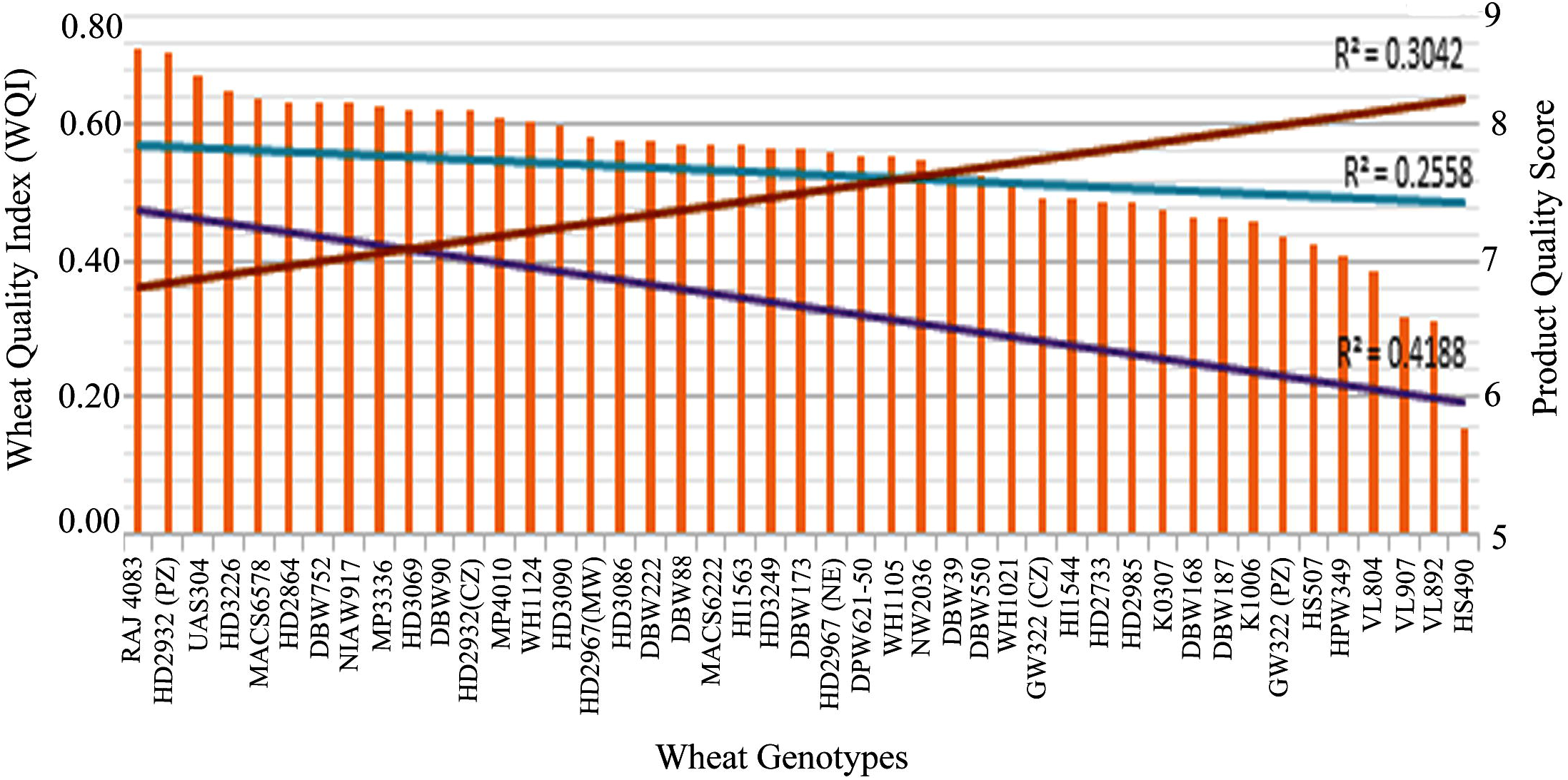
Wheat quality index (WQI) concerning quality of the end-products. X-axis represents the wheat genotypes used in the present study. Y-axis represents wheat quality index (left side) and product quality score (right side).

### 4.3 Application

It’s quite clear that wheat quality index makes a clear distinction in germplasm assortment. Breeders indeed need a single approach to discriminate the wheat lines for value addition but the quality of the end-products always dominate in the mind. Though wheat quality index provides reasonable surety in this endeavour, this system can be further refined to mark such preferences. All varieties in the elite group can be given score ‘A’ but the varieties with excellent *chapati* quality (score ≥ 8.0) or good bread quality (loaf volume ≥ 575cc and bread quality score ≥ 7.5) can be marked as A ^+^. If any varieties of the elite group excel in both the products, it can be glorified with A ^++^. Similarly, if a variety clubbed in the poor group show better product quality like cookies (HS 490 and HPW 349) or *chapati* (DBW 168), they can also be demarcated as C ^+^.

Besides germplasm assortment, this technique can also be immensely useful in the identification of the quality rich pockets or region. The usefulness of this strategy has been well demonstrated under Indian conditions where quality rich pockets have been identified in a high productivity zone *i.e*. NWPZ. Value addition characteristics of a zone or production environments have also been exemplified in this investigation.

## 5. Conclusion

Distinction for quality attributes is very important in wheat and the present system to rank superiority on multiple traits basis is laborious and demanding. This arduous method of screening for each quality trait can be done away by this suggested holistic approach where a single parameter *i.e*. wheat quality index is sufficient to sort out rank superiority in germplasm screening or identify the pockets suitable to harness the desired grain quality. It is further suggested to utilize this index in the screening of large number of breeding populations at both segregating and non-segregating generations. Since industry demands product specific wheat varieties to obtain superior product quality, this holistic approach could be a method to identify product-specific genotypes based on multi-traits. This also has implications on wheat growers by getting remunerative price to the growers through segregated procurement.

## Acknowledgements

The work is an outcome of a core project funded by ICAR (Project No.CRSCIIWBRCIL201500100182), New Delhi and the authors express their sincere thanks to the Director, ICAR-IIWBR for permitting the use of the data generated in the All India Coordinated Research Project (AICRP) on Wheat and Barley for this analysis. The efforts made by associated wheat research workers in sending grain samples of quality analysis and data reporting are also acknowledged.

## Authors’ contribution

Conceptualization (DM); Designing of the experiments (DM, GPS); Data compilation (GK, RS, OPG, VP); Formal analysis and interpretation (DM, RS, GK); Preparation of original draft (DM, RS, VP); Revision of manuscript (DM, OPG, GPS).

